# diffGEK: Differential Gene Expression Kinetics

**DOI:** 10.1101/2024.08.21.608952

**Authors:** Melania Barile, Shirom Chabra, Tomoya Isobe, Berthold Gottgens

**Author notes:** These authors contributed equally (Melania Barile, Shirom Chabra).

## Abstract

A defining characteristic of all metazoan organisms is the existence of different cell states or cell types, driven by changes in gene expression kinetics, principally transcription, splicing and degradation rates. The RNA velocity framework utilizes both spliced and unspliced reads in single cell mRNA preparations to predict future cellular states and estimate transcriptional kinetics. However, current models assume either constant kinetic rates, rates equal for all genes, or rates completely independent of progression through differentiation. Consequently, current models for rate estimation are either underparametrised or overparametrised. Here we developed a new method (diffGEK) which overcomes this issue, and allows comparison of transcriptional rates across different biological conditions. diffGEK assumes that rates can vary over a trajectory, but are smooth functions of the differentiation process. Analysing Jak2 V617F mutant versus wild type mice for erythropoiesis, and Ezh2 KO versus wild type mice in myelopoiesis, revealed which genes show altered transcription, splicing or degradation rates between different conditions. Moreover, we observed that, for some genes, compensatory changes between different rates can result in comparable overall mRNA levels, thereby masking highly dynamic changes in gene expression kinetics in conventional expression analysis. Collectively, we report a robust pipeline for comparative expression analysis based on altered transcriptional kinetics to discover mechanistic differences missed by conventional approaches, with broad applicability across any biomedical research question where single cell expression data are available for both wild type and treatment/mutant conditions.

## Introduction

There is no doubt that single-cell trancriptomic profiling represents a transformative technique to interrogate differentiation processes. By providing a snapshot of the RNA expression on an individual cell level, across a population of thousands of cells, each at distinct points in the dynamic process under study, differentiation trajectories can be reconstructed. These trajectories are inherently dynamic, characterised by changes in both the programmes and magnitudes of gene expression, as cells progress through a repertoire of cell states. Importantly, varying conditions - whether cell intrinsic, such as genetic mutations, or cell extrinsic, such as inflammation or pharmacological manipulation - can induce altered differentiation processes. Understanding the biology of how distinct conditions drive differing trajectories is crucial for understanding disease pathogenesis, drug mechanisms and stem cell engineering protocols.

The high resolution of scRNA-seq, combined with the wealth of transcriptomic information it provides, presents an opportunity to disentangle heterogeneity in differentiation processes between conditions, and unravel the mechanistic underpinnings that drive these differences. Differential gene expression (DGE) remains the predominant method to bridge the gap between observable phenotypes and the corresponding molecular mechanisms, evidenced by DGE often being the final (or near final) step in scRNA-seq analysis pipelines. A key problem, however, is that the majority of statistical or computational analysis procedures to achieve this either (i) fail to exploit the continuous resolution provided by scRNA-seq trajectories, or (ii) focus exclusively on the abundance of spliced mRNA, which limits interpretability of the underlying biological changes.

In fact, despite the continuous nature of differentiation trajectories, differential gene expression techniques predominantly adopt a discrete approach. More specifically, gene expression is often compared on discrete groups of cells in the differentiation pathway, e.g., by comparing clusters of cells by means of T-test based methods or upon modelling gene expression variance as a function of expression average (Love et al., 2014; Robinson et al., 2009). Critically these discrete approaches fail to account for the continuous dynamics of gene expression changes. Moreover, the comparison of discrete cell clusters within or between lineages introduces interpretative challenges. In particular, ambiguity arises in determining: which clusters should be compared, the proper aggregation of results from various pairwise cluster comparisons at different trajectory points, or how to account for the non-independence of these comparisons.

A few methods, such as tradeSeq, GPfates and Monocle 2 (Van den Berge et al., 2020; Lönnberg et al., 2017; Qiu et al., 2017), have been developed to improve trajectory-based differential expression analysis by adopting a dynamic approach. These methods process the continuous gene expression profiles obtained by ordering cells according to their transcriptomics difference from a pre-defined root cell (pseudotime), and compare these profiles across conditions, on a gene-by-gene level to determine differential expression. While these techniques better account for the dynamic nature of differentiation, a caveat is that they only rely on the abundance of spliced mRNA transcripts. This provides a surface level view of transcriptomic state, obscuring interpretability into the changes in underlying transcription, splicing and degradation rates (TSDr) which govern the process of gene expression. Moreover, since gene expression is an emerging property of these rates, discrepancies among conditions may exist at the rate level without precipitating as variations in spliced mRNA abundance; such differences would be ”missed” when comparing conditions using differential gene expression. Hence, methods to infer TSDr profiles within trajectories and compare these between conditions are vital to further advance the field.

To derive TSDr from scRNA-seq data from common scRNA-seq protocols, without the need for supplementary experiments or alterations to sequencing methodologies, we leveraged the combination of spliced and unspliced transcript abundance and build upon the foundations of the RNA velocity framework. Up until now, the relative abundance of nascent (unspliced) and mature (spliced) transcripts have been utilised as a powerful feature to interrogate the future state of a cell and the pseudotemporal order of differentiation in a lineage. These analyses are rooted in the widely-recognised RNA velocity framework (La Manno et al., 2018; Bergen et al., 2020). The core premise of RNA velocity is that spliced and unspliced count dynamics are coupled and governed by three kinetic parameters: transcription, splicing, and degradation rates. While effective in simple systems, the original framework falls short in certain scenarios. For example, if the nascent transcript is noisy, analysis is affected and several autoencoder-based methods have been suggested to infer a common latent space that reduces noise and allows correct trajectory inference (Qiao and Huang, 2021). An other work has suggested to use a per gene latent time (Gao et al., 2022) to account for gene specific dynamics. Still, these methods assume that the kinetic rates remain fixed across the differentiation trajectory (with the possibility to switch on/off the trascription), and thus do not account for conspicuous kinetics change along a trajectory, typical of development and particularly erythropoiesis (Barile et al., 2021). To address this limitation, we advocate for the incorporation of cell-specific values for these rates, similarly to what has been proposed earlier (Li et al., 2023; Qiu et al., 2022), but with further constrains to reduce the huge dimensionality of the respective unknown parameter space.

This paper introduces diffGEK, a method for analysing trajectory-based differential gene expression kinetics in scRNA-seq data. By extending the RNA velocity framework, diffGEK initially estimates per-cell and per-gene kinetic parameters using known lineage and pseudotemporal ordering of cells for a specific condition. Additionally, diffGEK integrates a statistical strategy to discern whether a gene exhibits differential kinetics between any two biological conditions, across all possible permutations (Table 1). Evaluation of the pipeline on simulated data and its application to two murine haematopoietic stem cell differentiation datasets demonstrates diffGEK’s utility as a complementary tool to differential gene expression analysis. diffGEK introduces an additional perspective for unraveling heterogeneity between states, enhancing interpretability and fostering a more nuanced understanding of the underlying biology.

**Table 1:** Schematic of the eight possible models used to fit the data. ”same” means that the corresponding rate is forced to be the same across conditions. ”any” means that it can be different. *α*: transcription rate; *β*: splicing rate; *γ*: degradation rate; k: number of model parameters (eight spline nodes per rate, four initial conditions and four estimated errors).

| | $\alpha$ | $\beta$ | $\gamma$ | k |
| --- | --- | --- | --- | --- |
| M1 | same | same | same | 32 |
| M2 | any | same | same | 40 |
| M3 | same | any | same | 40 |
| M4 | same | same | any | 40 |
| M5 | any | any | same | 48 |
| M6 | any | same | any | 48 |
| M7 | same | any | any | 48 |
| M8 | any | any | any | 56 |

## Results

### Simulations

We performed systematic simulations to prove that our framework can classify genes whose kinetics are different among two conditions and to optimise the fitting and classification procedure (Figure 1). Since any of the three kinetic rates can be identical or different among the two conditions, we have in total 2^3^ = 8 possible configurations, which we refer to as models. For each of these eight models (M1-8, Table 1), we simulated one hundred gene dynamics according to the following steps. First, we generated per-cells values of the three kinetic rates (transcription, splicing and degradation), modelling them with eight-nodes natural cubic splines and in such a way to keep their values similar among nodes to avoid oscillating patterns (Figure 1A, Methods). Modelling the rates as functions of pseudotime accounts for the potential changes in kinetics along a trajectory. Then, we generated the values of the spliced and unspliced counts (Figure 1B) for each model and condition with the standard ODE system describing RNA velocity (La Manno et al., 2018; Bergen et al., 2020) at twenty predefined pseudotime points, but using the pre-generated cell-dependent rates. The corresponding eight hundred genes were refitted each with the eight possible models, and with three different values of a penalization parameter, (*ρ*, Figure 1C and 1D), which ensures that the splines are not wobbly, for a total of 19200 fits. Successively, we ranked the models per each gene according to their corrected Akaike Information Criterion (AICc, Figure 1C), which accounts for the number of unknown variables and the square sum of residuals, as weighted by an extra regularization term, *µ*, which was allowed to take three different values (Figure D and Methods). Depending on the values of the regularization terms, our procedure succeeds or fails to classify the ground truth (see for example Table 2). In general, we saw that it is very easy to classify M1 genes as those genes that were generated with Model 1, i.e., assuming no differential kinetics. It was also generally easy to correctly classify those genes for which only one kinetic parameter changes. It was more complicated, though, to ensure that genes with more than two parameters changing among conditions were not classified as genes with only one parameter changing. We thus scored the performance of our procedure based on how many rates were correctly classified with respect to the ground truth (Table 3 and Methods). In total there are four possible scenarios for the rates: True Positive (TP), False Negative (FN), False Positive (FP) and True Negative (TN). Figure 1D shows the histograms of TP, TN, FP and FN classifications for each combination of regularization parameters. We picked the combination *ρ* = 10 and *µ* = 0.01 to ensure a large number of TP and TN, and a small number of FP, even if this means trading a large number of FN. Once picked the regularization parameters, we have automatically the best prediction for the model used to generate each gene. Overall, we created a pipeline that can infer genes that have differential kinetics among two conditions and dubbed it diffGEK.

**Table 2:**
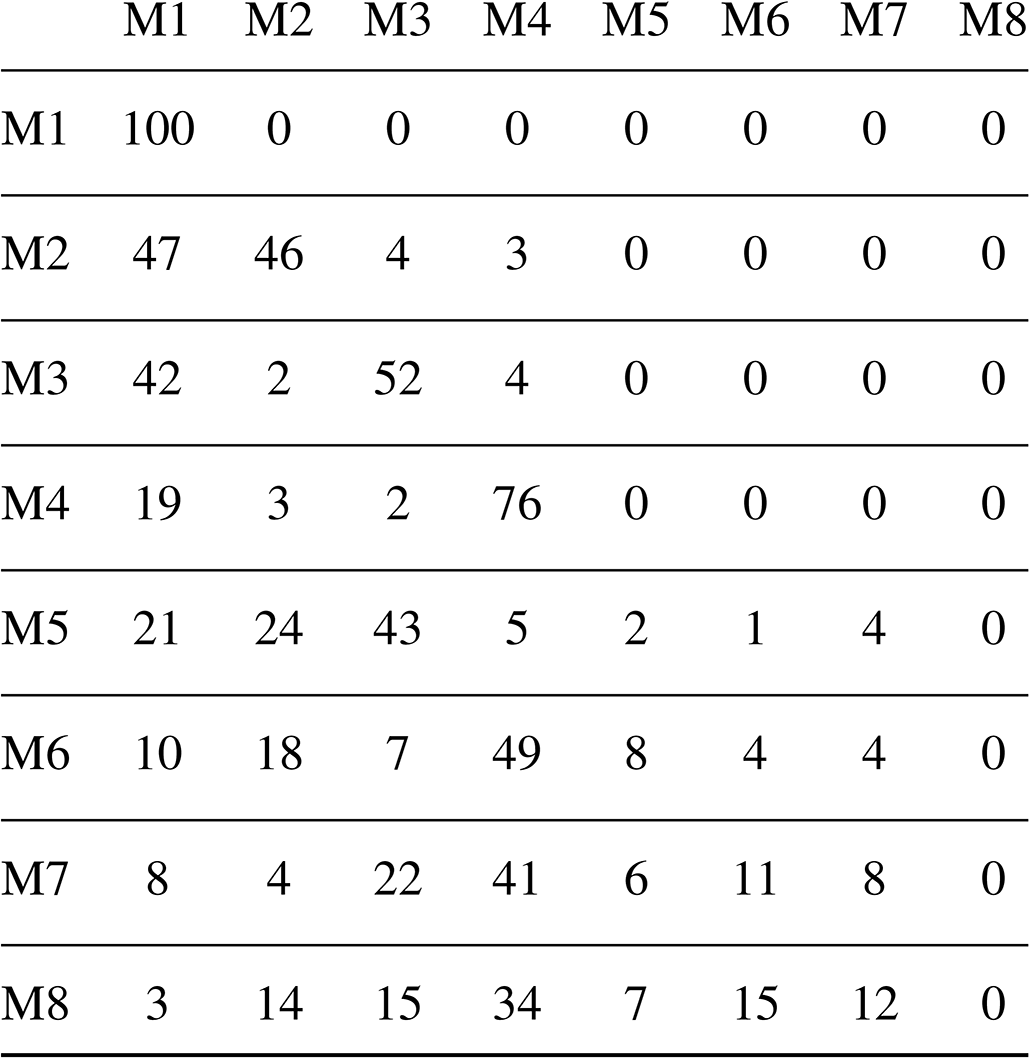
Example of classification of the 100 genes generated with each of the models on the rows, then refitted with the models in the columns, for a fixed set of regulatory terms. Perfect classification or non-significative classification is achieved in most cases where at most one parameter change among conditions, but it is more difficult when more than one parameter change. It is so needed to tune the regulatory terms to achieve the best deal among false/true positive/negative.

|  | M1 | M2 | M3 | M4 | M5 | M6 | M7 | M8 |
| --- | --- | --- | --- | --- | --- | --- | --- | --- |
| M1 | 100 | 0 | 0 | 0 | 0 | 0 | 0 | 0 |
| M2 | 47 | 46 | 4 | 3 | 0 | 0 | 0 | 0 |
| M3 | 42 | 2 | 52 | 4 | 0 | 0 | 0 | 0 |
| M4 | 19 | 3 | 2 | 76 | 0 | 0 | 0 | 0 |
| M5 | 21 | 24 | 43 | 5 | 2 | 1 | 4 | 0 |
| M6 | 10 | 18 | 7 | 49 | 8 | 4 | 4 | 0 |
| M7 | 8 | 4 | 22 | 41 | 6 | 11 | 8 | 0 |
| M8 | 3 | 14 | 15 | 34 | 7 | 15 | 12 | 0 |

**Table 3:** Numbers of True Negative (TN), True Positive (TP), False Negative (FN) and False Positive (FP) rates. A gene generated with Models M1-8 (rows) is fitted with models M1-8 (columns). Depending on which model is picked as best model, there can be either 0, 1, 2 or 3 TN, TP, FP, FN rates (clockwise starting from top left in the group of 4 numbers at the intersection of each pair model used for generating vs model used for fitting).

|  | M1 | M2 | M3 | M4 | M5 | M6 | M7 | M8 |
| --- | --- | --- | --- | --- | --- | --- | --- | --- |
| M1 | 3 0<br>0 0 | 2 0<br>0 1 | 2 0<br>0 1 | 2 0<br>0 1 | 1 0<br>0 2 | 1 0<br>0 2 | 1 0<br>0 2 | 0 0<br>0 3 |
| M2 | 2 0<br>1 0 | 2 1<br>0 0 | 1 0<br>1 1 | 1 0<br>1 1 | 1 1<br>0 1 | 1 1<br>0 1 | 0 0<br>1 2 | 0 1<br>0 2 |
| M3 | 2 0<br>1 0 | 1 0<br>1 1 | 2 1<br>0 0 | 1 0<br>1 1 | 1 1<br>0 1 | 0 0<br>1 2 | 1 1<br>0 1 | 0 1<br>0 2 |
| M4 | 2 0<br>1 0 | 1 0<br>1 1 | 1 0<br>1 1 | 2 1<br>0 0 | 1 1<br>0 1 | 1 1<br>0 1 | 0 0<br>1 2 | 0 1<br>0 2 |
| M5 | 1 0<br>2 0 | 1 1<br>1 0 | 1 1<br>1 0 | 0 0<br>2 1 | 1 1<br>0 1 | 0 1<br>1 1 | 0 1<br>1 1 | 0 2<br>0 1 |
| M6 | 1 0<br>2 0 | 1 1<br>1 0 | 0 0<br>2 1 | 1 1<br>1 0 | 0 1<br>1 1 | 1 2<br>0 0 | 0 1<br>1 1 | 0 2<br>0 1 |
| M7 | 1 0<br>2 0 | 0 0<br>2 1 | 1 1<br>1 0 | 1 1<br>1 0 | 0 1<br>1 1 | 0 1<br>1 1 | 1 2<br>0 0 | 0 2<br>0 1 |
| M8 | 0 0<br>3 0 | 1 0<br>2 0 | 1 0<br>2 0 | 1 0<br>2 0 | 0 2<br>1 0 | 0 2<br>1 0 | 0 2<br>1 0 | 0 3<br>0 0 |

**Figure 1.**
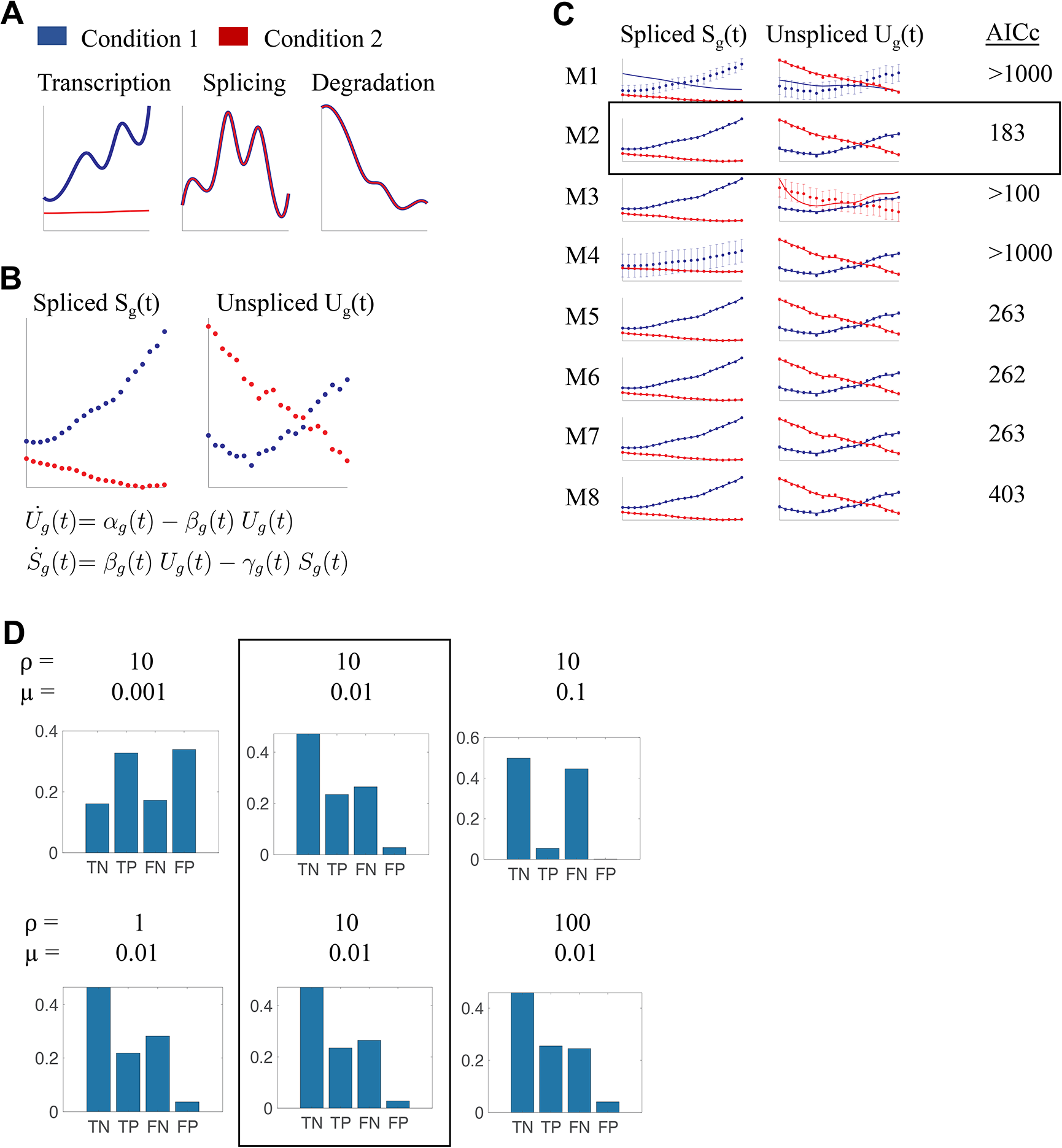
Schematic of diffGEK and the simulations performed to optimise it. (A) Generating the kinetic rates. The panel show an example of rates generated for two conditions (in blue and red) for a gene regulated according to Model 2 (different transcription rate, but same splicing and degradation rates). (B) Generating spliced and unspliced counts. Once the kinetic parameters have been generated, the unspliced and spliced counts are computed with the usual RNA velocity equations. (C) Each gene is refitted with all eight possible models. Dots: generated errors. Continuous lines: model fit. Error bars: error estimated by the fitting procedure. Then, the model corresponding to the smallest corrected Akaike Information Criteria index (AICc) is selected as best model for the corresponding gene and regularisation parameters (black rectangle). (D) For each pair of regularization parameters, we compute the number of rates that are correctly/incorrectly predicted. TN: True Negative; TP: True positive, FN: False Negative, FP: False Positive. The pair that gives the best compromise between TN, TP, FP, FN is selected (black rectangle). The diffGEK pipeline is ready.

### Transcriptional kinetics in Jak2 V617F mutant mice erythropoiesis

To evaluate diffGEK on biological data, we first applied the pipeline to a scRNA-seq dataset encompassing adult murine haematopoiesis from paired wild-type and homozygous Jak2 V617F mutant mice (Isobe et al., 2023). Homozygous Jak2 V617F mutations induces alterations in erythroid differentiation, resulting in a higher accumulation of mature erythroid cells in comparison to the paired wild-type controls (Figure 2A). Based on this understanding, we narrowed down our analysis to cells belonging to the erythropoietic trajectory, as determined by the cell fate probabilities computed by the original authors. Overall, the erythroid trajectory input into diffGEK consisted of 24,154 cells, derived from three sets of paired mutant and wild type mice. Of note, this dataset was deeply sequenced, with a median of 14,389 transcripts per cell, enabling reliable capture of unspliced reads. To input into the diffGEK pipeline, we integrated the top 1000 highly variable genes (HVGs) alongside the 80 genes identified as differentially expressed along the trajectory via tradeSeq, resulting in a collective set of 1042 genes due to list overlap (Figure 2B and Methods). Among these genes, 55 exhibited differential kinetics: 41 differed in transcription, 12 in splicing, and 2 in degradation rates (Figure 2C). In this Jak2 V617F mutation context, genes demonstrating differential gene expression kinetics displayed alterations in a single parameter only, rather than combinatorial parameter changes. Hyperge-ometric test confirmed the significant enrichment of genes with differential TSDr among those previously identified as differentially expressed genes through tradeSeq (*p <<* 0.001). For such genes exhibiting differential expression alongside changes in gene expression kinetics, diffGEK offers explanatory insights into the kinetic parameters driving the altered gene expression profile in the mutant (Figure 2D). Furthermore, since diffGEK adopts a continuous approach that is rooted in pseudotime, the output can be used to pinpoint the stages of differentiation, and the corresponding cell types, where kinetic rate differences are most pronounced (Figure 2E).

**Figure 2.**
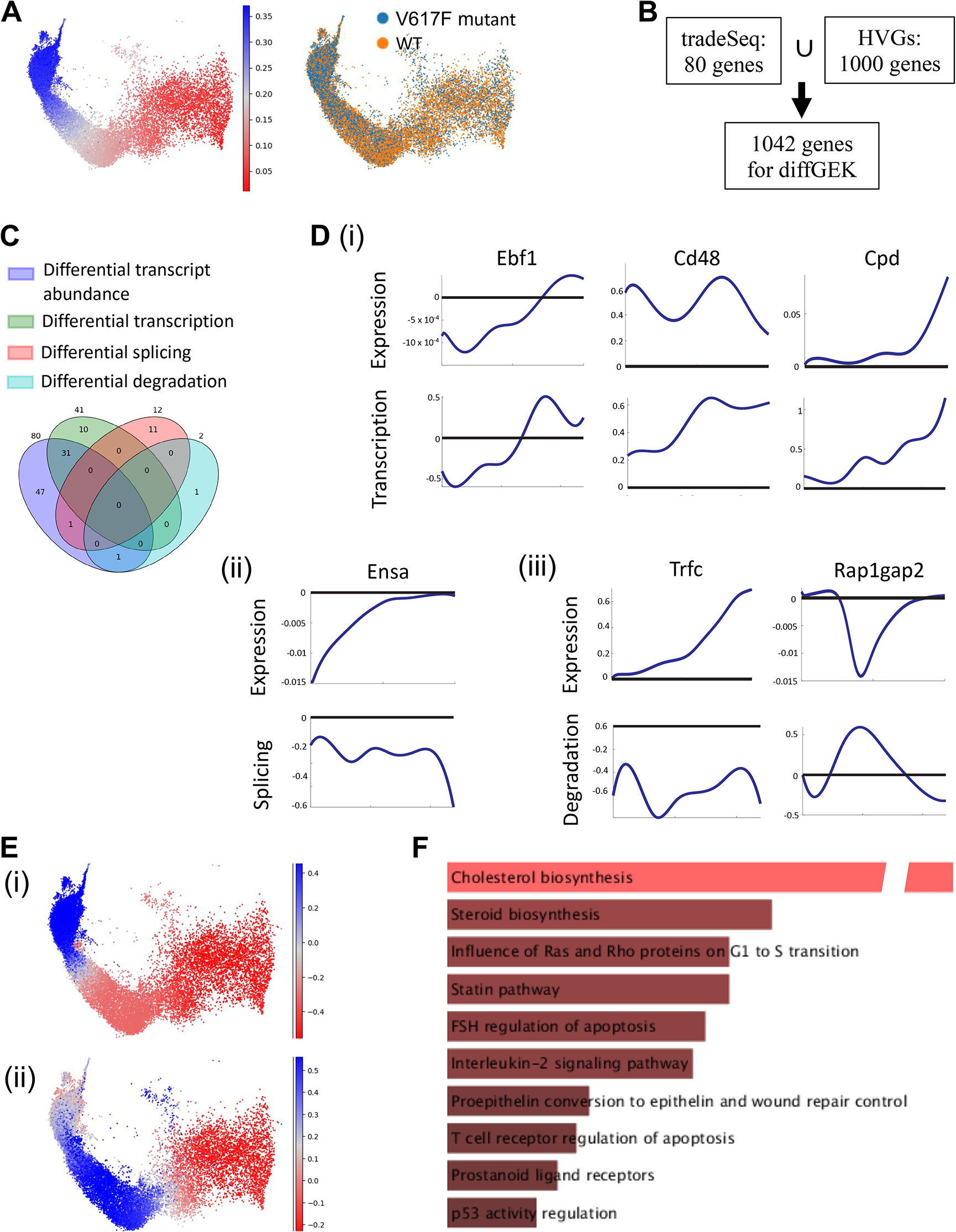
Application of the diffGEK pipeline to erythropoiesis in Jak2 V617F mutant vs WT mice. (A) UMAP showing diffusion pseudotime and conditions as adapted from the original manuscript (Isobe et al., 2023). (B) Choice of genes for the diffGEK pipeline. (C) Overlap between the differentially expressed genes and the diffGEK significant genes, sorted by rate type. (D) (i-iii) Examples of genes with differential kinetic rates (transcription, splicing or degradation respectively). For each gene, the panels represent the log2 fold change between the mutant and the wild type expression (top) and the rates (bottom). (E) Example of the log2 fold change of the transcription rate of Ebf1 (i) and the degradation rate of Rap1gap2 (ii). (F) Gene ontology of the diffGEK significant genes. Source: EnrichR, BioPlanet gene set. Adjusted pvalues *<* 0.01.

To gain further biological insight, we explored the gene ontology of genes displaying differential TSDr. Pathway enrichment analysis of these genes revealed a substantial overlap with pathways independently enriched from differentially expressed genes (Figure 2F). This highlights that diffGEK captures relevant differences between biological states, aligning with observations made by widely-utilised techniques. Key pathways exhibiting high enrichment scores included: (i) Cholesterol Biosynthesis and Steroid Biosynthesis, (ii) the p53 pathway, and (iii) the G1 to S transition. Notably, these pathways have been reported to play regulatory roles in erythroid differentiation, and were also enriched amongst differentially expressed genes. Cholesterol and steroid metabolism have recently been reported to control the balance between erythroid proliferation and differentiation, by regulating the cell cycle and induction of globin gene expression (Lu et al., 2022). These metabolic pathways are involved specifically during terminal erythroid differentiation which corroborates with the stage of the altered phenotype seen in the Jak2 V617F dataset. The p53 pathway operates comparably, controlling the switch between proliferation and differentiation, in addition to coordinating GATA1 expression in erythrocytes (Le Goff et al., 2021). In fact, aberrant activation of the p53 pathway has been associated with anaemia in ribosomopathies (Le Goff et al., 2021). Conversely, in Jak2 V617F, an inverse scenario might be in effect, potentially driving increased terminal differentiation. Finally, the regulation of cell cycle speed and critically, the rate of transition to S phase has recently emerged as another regulatory mechanism governing erythroid differentiation and output (Socolovsky, 2022).

It is important to stress that diffGEK also recovered differences in genes that were ”missed” by differential expression. Some of these genes, such as Cdkn1a, Bbc3 and Dusp1 belong to the p53 pathway that was enriched amongst differentially expressed genes; and likewise alterations in the kinetics of Nfkb1a, Cdkn1a were identified that belong to the cell cycle G1/S transition pathway. In addition other genes belonged to pathways that were not picked up by gene ontology on differential expression and warrant further exploration such as the intracellular transduction machinery downstream of the TNFR2 pathway and the CD40L pathway, which are targets commonly implicated in lymphocyte differentiation (extra pathways not shown). Globally, genes belonging to the same pathway demonstrated concordant kinetic changes, evidenced by strong enrichment for terms when looking at all genes with changes in either transcription, splicing or degradation. For example, genes with altered transcription rates were overrepresented for cholesterol and steroid metabolism, as well as the G1/S transition. Conversely, genes with altered splicing rates were enriched in immune signalling related pathways (CD40L, TNFR2, NFKB, IL-1 signalling) and apoptotic terms. These consistent patterns amongst pathway members may imply that different pathways are subjected to alterations in different processes i.e. transcription, splicing, degradation.

More broadly, this analysis underscores the value of exploring gene expression dynamics through scRNA-seq as an adjunct layer of comparison, to capture nuances that might be missed when solely examining differences in spliced mRNA abundance, highlighting new genes belonging to existing candidate pathways or identifying new candidate pathways altogether.

### Transcriptional kinetics in Ezh2 KO myelopoiesis

Next, we applied our framework to another published scRNA-seq dataset which followed the combinatorial effects of two oncogenic mutations, *NRasG*12*D*^+^*^/^* and *Ezh*2*^ko/ko^* on driving perturbed haematopoiesis across three timepoints (T1, T2, and T3) (Liu et al., 2021). By T2, the mice had developed myeloproliferative neoplasm, which by T3 had progressed to acute myeloid leukaemia, characterised by arrested myeloid differentiation and the accumulation of immature myeloid cells. Based on this, we narrowed down our analysis to cells belonging to the myeloid trajectory (Figure 3A) and applied the diffGEK pipeline to interrogate differences between the wild-type and mutant state. Overall, the myeloid trajectory input into diffGEK consisted of 11984 cells, derived from two sets of paired mutant and wild type mice. Of note, this dataset was deeply sequenced, with a median of 3144 transcripts per cell, enabling reliable capture of unspliced reads. Given the presence of a time course, we selected genes expressed differentially on the trajectory via tradeSeq, comparing conditions one time point at a time, and for all timepoints together. Altogether, we ran diffGEK for 1228 genes, including the top 1000 HVGs (Figure 3B). Since the most pronounced alterations to myeloid differentiation manifested at T3, we explored the results of diffGEK on the wild-type and mutant trajectory at this timepoint.

**Figure 3.**
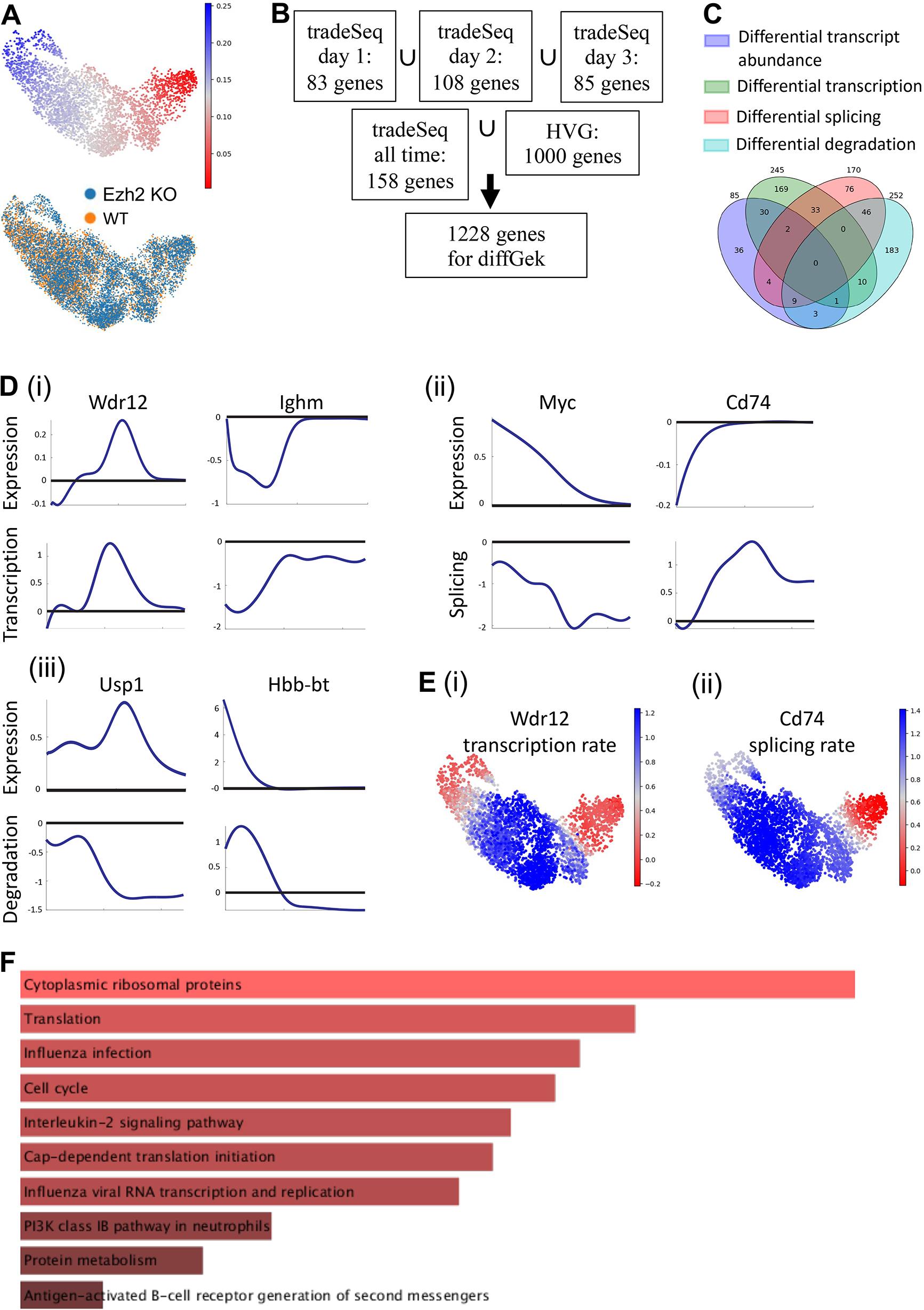
Application of the diffGEK pipeline to myelopoiesis in Ezh2 KO vs WT mice. (A) UMAP showing diffusion pseudotime and conditions as adapted from the original manuscript (Liu et al., 2021). (B) Choice of genes for the diffGEK pipeline. (C) Overlap between the differentially expressed genes and the diffGEK significant genes, sorted by rate type. (D) (i-iii) Examples of genes with differential kinetic rates (transcription, splicing or degradation respectively). For each gene, the panels represent the log2 fold change between the mutant and the

diffGEK recovered 667 genes with differential kinetics of which 428 genes had alterations in one parameter only, and 239 had changes in more than one parameter (Figure 3C). This highlights two key distinctions from the analysis of the pre-leukaemic Jak2 dataset: (i) there were globally a higher number of genes with differential kinetics, and (ii) there were genes with multiple parameter changes which may be aligned with increased dysregulation in the leukaemic state. Moreover, hypergeometric test again showed statistically significant overlap between genes identified to have distinct TSDr and altered transcript abundance (differentially expressed genes). For such genes that had altered TSDr and differential expression, diffGEK provides an interpretability layer to understand the kinetic process driving the altered transcript abundance, as seen by the examples in Figure 3D. As in the previous case, kinetic differences can be visualized along the differentiation trajectory, to pinpoint the cell types where they are more pronounced, which are gene-dependent (Figure 3E).

Next, we explored the gene ontology of those genes identified with differential TSDr (Figure 3F). It is important to first note that 85 differentially expressed genes identified by tradeSeq revealed statistically significant enrichment for two pathways only. In contrast, pathway enrichment analysis of genes identified by diffGEK demonstrated significant enrichment for a number of relevant terms. Pathways with the highest enrichment score, included: translation and ribosomal proteins, whose intricate regulation is perturbed in myeloid leukaemias and is a driver for protein metabolism that promotes elevated cell cycling which was another highly enriched term. In addition, other pathways with high enrichment consisted of interleukin-2 signalling, which has been reported to be a key signalling pathway required for the survival of leukaemia initiating cells in chronic myeloid leukaemia (Kobayashi et al., 2014); moreover this has been shown to be an early change. The Class IB PI3K pathways, and Aurora B pathways were also enriched and have published roles in driving AML (Darici et al., 2020), and AML t(8;21) (Qi et al., 2019) respectively.

Taken together, this analysis highlights that diffGEK is able to uncover changes in TSDr, early in transformation before they are picked up by differential expression, in important genes belonging to candidate pathways associated with myeloid leukaemia.

## Discussion

Here we present, diffGEK, a comparative technique which leverages both spliced and unspliced mRNA counts, to extract trajectory-based differential gene transcription, splicing and degradation rates between conditions. diffGEK builds on the current suite of scRNA-seq comparative tools by enabling researchers to factor in the continuous nature gene expression profiles inherent to differentiation trajectories, while diving deeper to deliver interpretability into the gene expression kinetics changes that drive altered transcriptional profiles. We propose for diffGEK to be applied to the end of bioinformatic pipelines as an adjunct to differential gene expression, where it can provide a more granular approach to extract differences between conditions which may not precipitate into altered mRNA abundance, while providing additional interpretability.

diffGEK derives TSDr from conventional scRNA-seq datasets, without the need for metabolic labelling experiments, or alterations to single cell sequencing protocols. This is achieved through leveraging unspliced reads, which are a largely underused modality in scRNA-seq. One of the major attractions of utilising the unspliced reads layer is that the added information provided is a low-hanging fruit which is essentially available at no additional cost as it is extracted from the raw scRNA-seq counts. It is worth noting that existing scRNA-seq technologies are not optimised for capturing intronic reads with high efficiency. This is due to their heavy reliance on priming methods, which sequence either the 3’ or 5’ ends of genes while neglecting the intervening regions. However, the detection of intronic reads can be plausibly attributed to secondary priming positions of oligo dT onto short stretches of poly-A sequences within introns. Despite not being optimised for the capture of intronic reads, scRNA-seq protocols are capable of recovering this modality in significant depth. In fact, an analysis of single-cell RNA-seq datasets derived from a range of leading protocols such as SMART-seq2, STRT/C1, inDrop and 10x Genomics Chromium found that 15–25% of all captured reads were unspliced sequences. We must acknowledge a caveat concerning scRNA-seq alignment frameworks, which primarily attribute the presence of introns as a consequence of incomplete processing of pre-mRNA. However, it is important to recognise that other mechanisms, such as intron retention, can also contribute to the presence of intronic sequences in fully processed cytoplasmic mRNA molecules. Consequently, these intronic sequences may be erroneously assigned as unspliced reads. However, to address this concern, La Manno et al. conducted metabolic labelling to show that the number of unspliced reads recovered by scRNA-seq correlate with metabolically labelled nascent mRNA molecules, and the level of scRNA-seq spliced counts, suggesting that these intron-containing molecules recovered by scRNA-seq predominantly represent unspliced precursor mRNAs; this emphasizes the considerable potential for the development of methodologies explicitly designed to enhance the retrieval of unspliced and intronic sequences. Moreover, the integration of approaches such as single-cell metabolic labelling, which provide ground-truth measurements of unspliced mRNA, could further enrich the pipeline and amplify the signal-to-noise ratio. It is important to note that our diffGEK analysis fully supports the notion that unspliced reads extracted even with current sequencing technologies and alignment methods, are sufficient to provide meaningful gene expression kinetic rates. These TSDr effectively explain differentially expressed genes and enrich for relevant biological pathways. This solidifies the inherent value of unspliced counts as genuine biological signal, thereby extending the usability of this analysis to the majority of scRNA-seq datasets.

At its core, diffGEK builds upon and refines the RNA velocity framework. The significant innovation introduced by RNA velocity, in modelling differentiation dynamics through the relative abundance of spliced and unspliced reads, has come with a number of technical issues. Ever since the seminal 2019 paper by La Manno et al., several subsequent studies have tackled its limitations. These advancements include accommodating varying gene activation states along a trajectory (as demonstrated in Bergen et al.), employing latent space representations to counteract inference errors due to count noise, especially within unspliced counts (Qiao and Huang, 2021), integrating cell-specific kinetic rates to accommodate kinetic enhancements (Barile et al., 2021), and addressing potential alterations in transcriptional kinetics within differentiating systems (as indicated in a recent publication). In this work, we particularly focused on comparing genes with kinetic changes (differential TSDr) among conditions. To this aim, we expanded upon the existing RNA velocity paradigm by enabling cell-specific kinetic rates within a predetermined trajectory based on precomputed pseudotemporal cell order. Contrary to the approach in cellDancer (Li et al., 2023), which assumes independent kinetic rates for each cell, our approach smoothed the rates over a trajectory. Despite being a strong assumption, the method proved to be significantly more stable, especially considering that in our test datasets, independent cell kinetics failed to recapitulate the known biology (data not shown).

diffGEK provides a flexible framework that can be used downstream of any dimensionality reduction and trajectory inference method. As it is agnostic to the dimensionality reduction and trajectory inference methodology, the approach scales from simple to complex trajectories with multiple bifurcations: diffGEK only requires the original spliced and unspliced expression count matrix of the individual cells, estimated pseudotimes, and a hard assignment of the cells to the lineages. The flexibility provided by diffGEK to work downstream of any trajectory inference technique is crucial, as trajectory-based DE is often the final (or near final) step in a much longer analysis pipeline. Nevertheless, it is worth mentioning that our method, while allowing for flexibility, as with other terminal scRNA-seq analytical tools still remains strongly dependant on the pre-defined trajectories and pseudotime. A different definition of trajectories and pseudotime could identify different kinetically variable genes. Furthermore, we relied on the assumption that the TSDr can be modelled with smoothed splines, differently from our previous attempt to model them with completely discontinuous transitions (Barile et al., 2021). The discontinuous transitions can still be obtained in our current framework upon increasing the number of spline nodes, but has a computational cost and an overfitting risk. Nevertheless, our pipeline has proven to bring insights in experiments with multiple conditions and allows for further improvement, such as unsupervised learning of trajectories and latent time, and higher computational efficiency.

Our tool has been optimised and coded for comparison of two conditions, but it can easily modified for multiple conditions, provided that the optimisation parameters are re-optimised similarly to what described here. Note, though, that the computational power scales with the number of compared conditions in a power-like fashion.

diffGEK offers a range of benefits for biological interpretation. It offers an explanatory lens into the root causes of differential transcript abundance observed in differential gene expression, providing a finer resolution to guide potential corrective strategies. In addition to enhancing interpretability, diffGEK probes beyond surface level spliced mRNA expression to uncover nuanced differences that may not be reflected in differential transcript abundance. In doing so this method can facilitate the identification of potential new gene candidates within enriched pathways or the discovery of entirely novel candidate pathways that could be instrumental in driving the observed altered phenotype. It is important to stress that the interpretability afforded by diffGEK operates on two levels. It not only clarifies the nature of kinetic alterations (transcription, splicing, and degradation) but also identifies specific pseudotime locations along the trajectory where substantial differences arise. This understanding has the potential to facilitate targeted therapeutic approaches directed at specific cell types and distinct processes of the mRNA lifecycle. Indeed a range of RNA-based and protein-based modulators have emerged which employ an array of strategies to modulate transcription, splicing and degradation rates. However, the majority of these modifiers lack gene-specific action, exerting their effects globally across multiple genes.We anticipate that diffGEK may provide valuable insights into the pathophysiology of various solid tumors and haematological malignancies carrying mutations in genes that modulate TSDr. Notably, mutations in spliceosomal genes such as SF3B1, U2AF1, SRSF2, ZRSR2, and the RNA-stabilizing methyltransferase METTL3 are specifically enriched in haematological disease. This is particularly pertinent in the haematological system as conventional analysis suites for differential gene expression or alternative splicing analysis have provided limited understanding of disease mechanisms. Nevertheless, our analysis of the two datasets presented in this study reveals that mutations in genes unrelated to transcription, splicing, and degradation can also influence TSDr. This highlights the broader applicability of diffGEK in exploring various cell-intrinsic mutational contexts, as well as cell-external factors like pharmacological drugs and cell-reprogramming cytokines, which drive diverse cellular states.

## Methods

### General approach

To model the spliced and unspliced counts over pseudotime we relied on the previously established coupled ODE system:

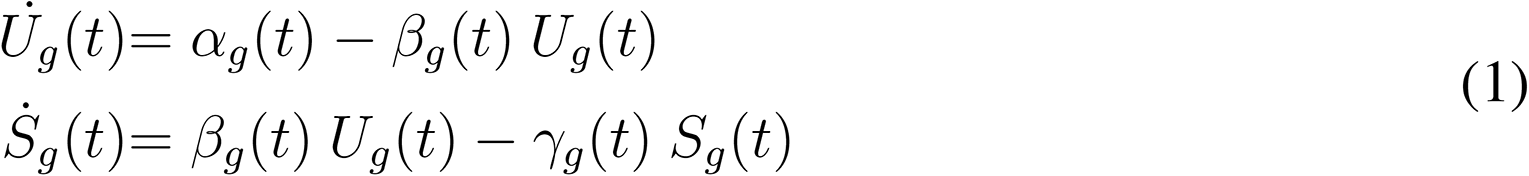

where *U_g_*(*t*) and *S_g_*(*t*) are the number of unspliced and spliced counts for gene *g* at pseudotime *t*, respectively; *α_g_*(*t*), *β_g_*(*t*) and *γ_g_*(*t*) are the kinetic parameters (transcription, splicing and degradation rate respectively) for gene *g* at pseudotime *t*. Compared to other papers, we modelled the kinetic parameters in a novel way. First, the parameters are gene specific. Second, they are a function of the cellular state, i.e., of the pseudotime. Since this would imply a big number of unknown variables to estimate, we further assumed that the parameters can be modelled with natural cubic splines, and required that such splines are as least wobbly as possible (see Optimisation).

### Optimisation

To model *α_g_*(*t*), *β_g_*(*t*) and *γ_g_*(*t*), we chose natural cubic splines with 8 nodes per each parameter. Furthermore we included 2 more variables to account for the initial conditions, and two more to account for the estimated error on each time course (spliced and unspliced counts over pseudotime). So in total we have 28 unknown variables per gene. We calibrated the model output to the simulated or real data pooled in 20 values of the pesudotime. So for each dataset, each pool contains a number of cells corresponding to the dataset size divided by 20 (floored value). The values used for fit are the averaged values of the spliced/unspliced counts, and the averaged values of the corresponding pseudotimes. For simulations, we generated the counts directly. For real data, we imputed the normalized counts with the standard scVelo pipeline. For each gene independently, we maximized the following negative loglikelihood function:

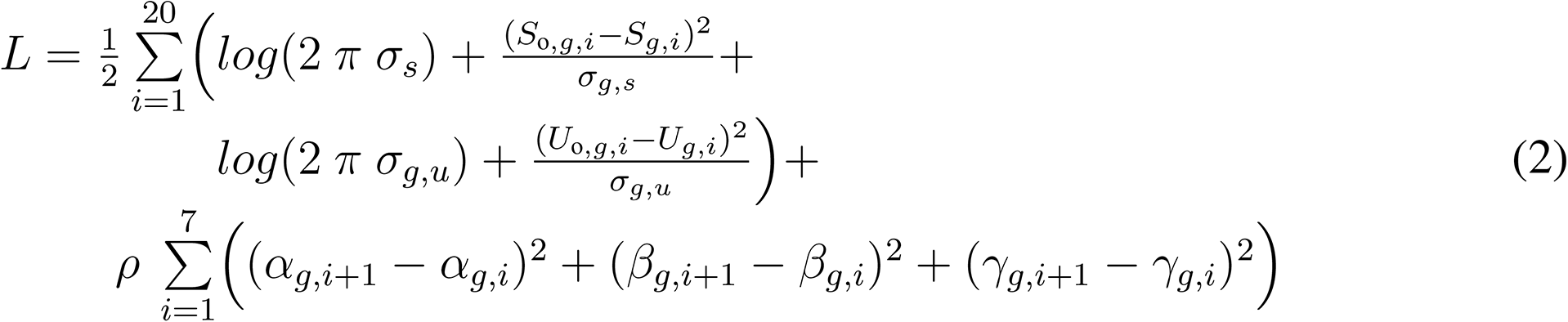

where *U*_o_*_,g,i_* and *S*_o_*_,g,i_* are the measured pooled values of the unspliced and spliced counts respectively for gene *g* at the pooled pseudotime value *i*; *U_g,i_* and *S_g,i_* are the predicted values of the unspliced and spliced counts respectively for gene *g* at the pooled pseudotime value *i*; *σ_g,s_* and *σ_g,u_* are the estimated variance on the all time course of the spliced and spliced counts for gene *g* respectively; *ρ* is a parameter (to be fixed with simulations) regularizing the wobbliness of the splines; for the kinetic rates, the sum runs over the splines nodes. The optimisation was performed in MATLAB 2019 with the toolboxes AMICI and PESTO. For each gene, we launched 3000 optimisations starting from randomly picked initial points. Constrains on parameters were as follows:

- kinetic parameters: from 0 to 6.7 per unit of pseudotime;
- estimated initial values: from 10% to 300% of the measured initial value;
- estimated errors: from 1% to 10% of the maximum value of the corresponding data.

### Model selection

The framework described so far allows the estimation of the kinetic parameters in units of the pseudotime. While this can be informative on the variation of the rates along a trajectory, it is also difficult to interpret given that the pseudotime unit is arbitrary. We can nevertheless gain information of what rates change among different conditions. For a comparison among two conditions, for example, there can be 8 possible scenarios (see Table 1). We fit these 8 models simultaneously to the spliced and unspliced counts for each condition; for each gene, each of the 8 models is optimised independently. After optimisation, for each genes we obtain the estimated parameters for each of the 8 models. In order to choose the best model, we first need to evaluate the goodness of the fit. We found empirically that the value of the negative loglikelihood is not good to compare different models, because the error per time course is estimated independently and can be very different across models (neither it helps to force such error to be the same, given that the values for the 2 conditions can be very different). We thus used a squared sum of residuals, *SSR_g,m_*, weighted by an extra parameter, *µ*, that has again to be determined via the simulations (see next section). We then ranked models according to their corrected Akaike index per gene *g* per model *m*, *A_g,m_*, which is calculated as follows: 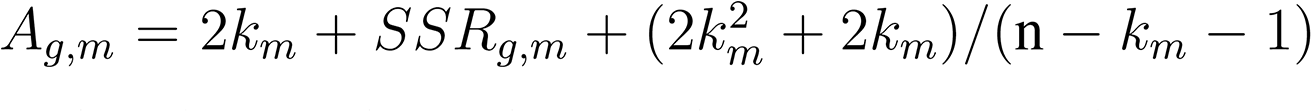, where *k_m_* is the number of parameters and n the fixed number of data (40 for all the example in this paper). The model with the minimum *A_g,m_* per each gene is picked as the best model.

### Simulations

In order to determine the values of *ρ* and *µ*, we simulated 100 genes for each of the 8 models described in the previous section . The genes are simulated with randomly extracted values of the initial conditions and of the first node of the parameters’ splines. The following nodes are extracted in a gaussian range of the previous node, to avoid massive wobbling in the parameters’ values. The corresponding 800 sets of 4 time courses (spliced and unspliced time courses for each of the two conditions) are re-fit, each with the 8 possible models, for a total of 1600 fits. For each gene simulated with model *m*, we would of course expect to recover model *m* as best model, while recovering a different model can imply having a certain number of true/false positive/negative outcomes. By false positive we mean any rate that is simulated as non changing but for which the selected model predicts a change among conditions, and so on (Box2). We found that the values for *ρ* = 10 and *µ* = 0.01 maximised the true positive and true negative while keeping the false positive and negative low (Figure 1E). We used this values for running our model on real data.

### Jak2

The Jak2 landscape was taken from Isobe et al., 2023. We selected the erythroid trajectory because we expect Jak2 to produce changes in this lineage. We ran diffGEK on the top 1000 HVGs as selected by the scVelo command *scv.pp.filter and normalize*, plus the tradeSeq differentially expressed genes as found by the authors in the original paper, for a total of 1048 genes.

### Ezh2

The Ezh2 dataset was taken from Liu et al., 2021, accession GSE179084. We downloaded 6 raw fastq data (GSM4711730, GSM4711731, GSM4711732, GSM4711739, GSM4711740, GSM4711741) corresponding to the 3 time points for WT and Ezh2-KO conditions; we ran cellranger and then velocyto to obtain spliced/unspliced counts. We then created an UMAP following the standard scanpy pipeline, and identified myeloid progenitors with celllrank. We then ran tradeSeq to get differential trajectories for the genes at all time points together and for each time point separately. We ran diffGEK on all such genes plus 1000 top HVGs, for a total of 1228 genes.

### Ontology analysis

For both Jak2 and Ezh2 we performed ontology analysis on the EnrichR website. For both cases, we used the genes that have differential kinetics, and took the BioPlanet gene set. The adjusted maximum pvalues are specified in the respective figure legends.

### Code and data availability

For this study we did not produce new data. Jak2 data was taken from Isobe et al., 2023. Ezh2 data was taken from Liu et al., 2021. Simulated data are available on request. All the codes have been uploaded to GitHub: https://github.com/mebarile/transcriptional_kinetics.

### Authors contribution

M.B., S.C. and B.G. conceived the study. M.B. and S.C. designed the pipeline and the downstream analysis. M.B wrote and ran the codes, S.C. analysed and interpreted the output. T.I. supervised and contributed to the analysis of the erythropoietic dataset. M.B., S.C. and B.G. wrote the manuscript, and T.I. proofread and edited it.

## Acknowledgments

Work in the Gottgens laboratory is supported by Wellcome (206328/Z/17/Z and 203151/Z/16/Z), Blood Cancer UK(18002), Cancer Research UK (C1163/A21762), and UKRI Medical Research Council (MC PC 17230). M.B. was further funded by the Centre of Translational Stem Cell Biology, Hong Kong SAR, China.

## Supplementary note

### Comparison with rates estimated by different methods

diffGEK is a tool for estimating kinetics rates given a trajectory. It does not attempt to estimate a cellular ordering. This makes the comparison with other tools that estimate the cellular ordering difficult: indeed, we do not know if differences arise because of the methods that estimates the rates, or because the difference in the estimated/assumed cellular ordering. Nevertheless, we tested two methods (scVelo, from Bergen et al., and cellDancer from Li et al.) on our two biological datasets and compared the estimated rates among them and with diffGEK. Note that we cannot test the methods on the in silico datasets because we did not simulate the transcriptomic abundances, but we directly simulated the gene dynamics. We tested these methods separately on the WT and mutant/KO conditions, in keeping with diffGEK, whose pipeline is run on each condition independently. First of all, neither method could recapitulate the known biology with respect to the cellular ordering (Figure 4). The arrows, which represent the trajectories, should point from the stem cells to the erythroid/myeloid progenitors, but they actually point in different direction, if not completely backward. This is expected to significantly affect the rate estimates. We then compared the rates for each pair of methods. Note, however, that even this comparison is problematic. Indeed, scvelo recovers a set of 3 rates per gene, because the rates are assumed to be constant along a trajectory. cellDancer and diffGEK recover a rate per gene and per cell. So, in order to compare the latter with scvelo, we took the average rate per gene and then computed the pearson correlation. To compare cellDancer and diffGEK among them, we computed the correlation considering each corresponding rate per gene per cell. Of note, the rates are in units of pseudotime/latent time, which has different scales depending on the method; still, we used the correlation coefficient because we assumed that the units of measurement are scaled linearly. The results are shown in Tables 4-7. It is obvious that the comparison is very poor, with most correlation coefficients being close to zero. The only rates that seemed to be correlated are the transcription rates between diffGEK and scvelo, while cellDancer does not correlate for any rate. As stated above, however, the methods are very different and the correlations itself may not be very informative on the performance.

**Table 4:**
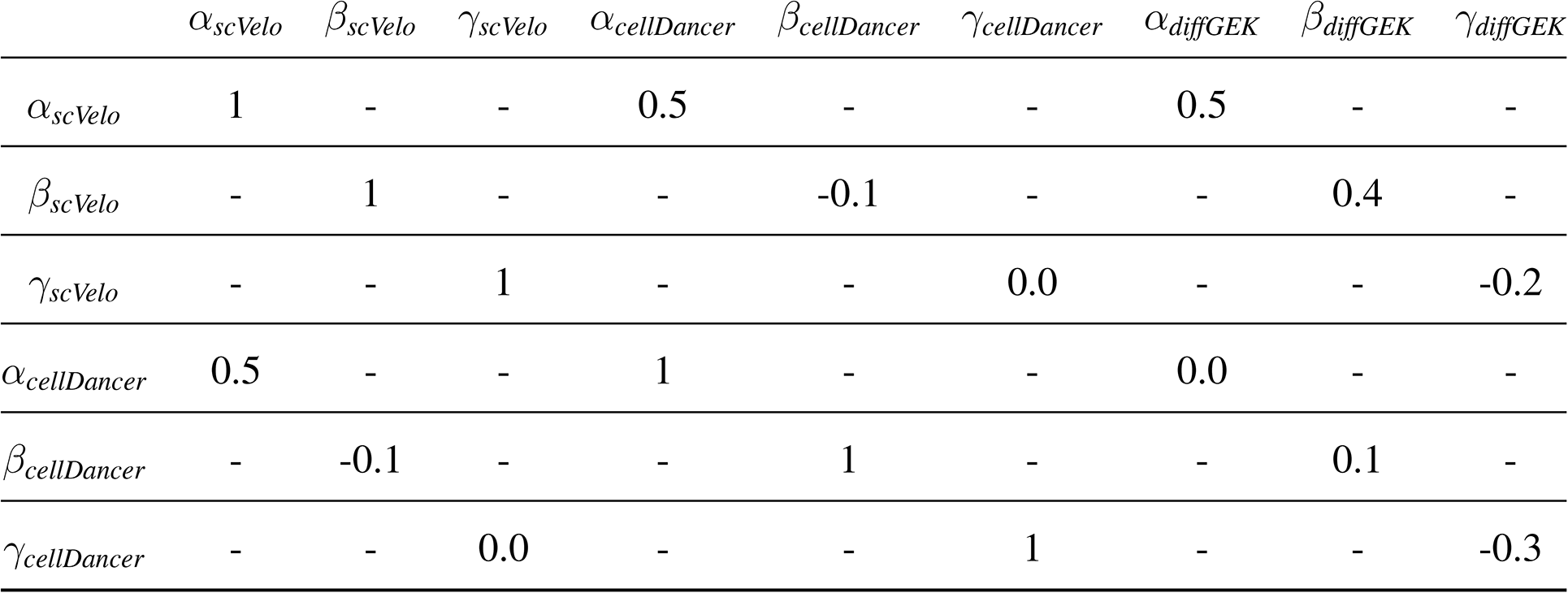
Methods comparison for erythropoiesis in WT mice. *alpha*: transcription rate; *beta*: splicing rate; *gamma*: degradation rate. Numbers represent the Pearson correlation coefficient of the estimated parameters as computed with the corresponding tool (see Methods).

**Table 5:** Methods comparison for erythropoiesis in Jak2 V617F mutant mice. *alpha*: transcription rate; *beta*: splicing rate; *gamma*: degradation rate. Numbers represent the Pearson correlation coefficient of the estimated parameters as obtained with the corresponding tool (see Methods).

| | $\alpha_{scVelo}$ | $\beta_{scVelo}$ | $\gamma_{scVelo}$ | $\alpha_{cellDancer}$ | $\beta_{cellDancer}$ | $\gamma_{cellDancer}$ | $\alpha_{diffGEK}$ | $\beta_{diffGEK}$ | $\gamma_{diffGEK}$ |
| --- | --- | --- | --- | --- | --- | --- | --- | --- | --- |
| $\alpha_{scVelo}$ | 1 | - | - | 0.1 | - | - | 0.5 | - | - |
| $\beta_{scVelo}$ | - | 1 | - | - | -0.1 | - | - | 0.4 | - |
| $\gamma_{scVelo}$ | - | - | 1 | - | - | 0.0 | - | - | -0.2 |
| $\alpha_{cellDancer}$ | 0.1 | - | - | 1 | - | - | 0.0 | - | - |
| $\beta_{cellDancer}$ | - | -0.1 | - | - | 1 | - | - | 0.0 | - |
| $\gamma_{cellDancer}$ | - | - | 0.0 | - | - | 1 | - | - | -0.4 |

**Table 6:** Methods comparison for myelopoiesis in WT mice. *alpha*: transcription rate; *beta*: splicing rate; *gamma*: degradation rate. Numbers represent the Pearson correlation coefficient of the estimated parameters as obtained with the corresponding tool (see Methods).

| | $\alpha_{scVelo}$ | $\beta_{scVelo}$ | $\gamma_{scVelo}$ | $\alpha_{cellDancer}$ | $\beta_{cellDancer}$ | $\gamma_{cellDancer}$ | $\alpha_{diffGEK}$ | $\beta_{diffGEK}$ | $\gamma_{diffGEK}$ |
| --- | --- | --- | --- | --- | --- | --- | --- | --- | --- |
| $\alpha_{scVelo}$ | 1 | - | - | 0.3 | - | - | 0.6 | - | - |
| $\beta_{scVelo}$ | - | 1 | - | - | 0.0 | - | - | 0.2 | - |
| $\gamma_{scVelo}$ | - | - | 1 | - | - | 0.1 | - | - | -0.1 |
| $\alpha_{cellDancer}$ | 0.3 | - | - | 1 | - | - | 0.0 | - | - |
| $\beta_{cellDancer}$ | - | 0.0 | - | - | 1 | - | - | 0.1 | - |
| $\gamma_{cellDancer}$ | - | - | 0.1 | - | - | 1 | - | - | -0.1 |

**Table 7:** Methods comparison for myelopoiesis in Ezh2 KO mice. *alpha*: transcription rate; *beta*: splicing rate; *gamma*: degradation rate. Numbers represent the Pearson correlation coefficient of the estimated parameters as obtained with the corresponding tool (see Methods).

| | $\alpha_{scVelo}$ | $\beta_{scVelo}$ | $\gamma_{scVelo}$ | $\alpha_{cellDancer}$ | $\beta_{cellDancer}$ | $\gamma_{cellDancer}$ | $\alpha_{diffGEK}$ | $\beta_{diffGEK}$ | $\gamma_{diffGEK}$ |
| --- | --- | --- | --- | --- | --- | --- | --- | --- | --- |
| $\alpha_{scVelo}$ | 1 | - | - | 0.2 | - | - | 0.4 | - | - |
| $\beta_{scVelo}$ | - | 1 | - | - | 0.0 | - | - | 0.2 | - |
| $\gamma_{scVelo}$ | - | - | 1 | - | - | 0.2 | - | - | 0.0 |
| $\alpha_{cellDancer}$ | 0.2 | - | - | 1 | - | - | 0.0 | - | - |
| $\beta_{cellDancer}$ | - | 0.0 | - | - | 1 | - | - | 0.0 | - |
| $\gamma_{cellDancer}$ | - | - | 0.2 | - | - | 1 | - | - | -0.1 |

**Figure 4.**
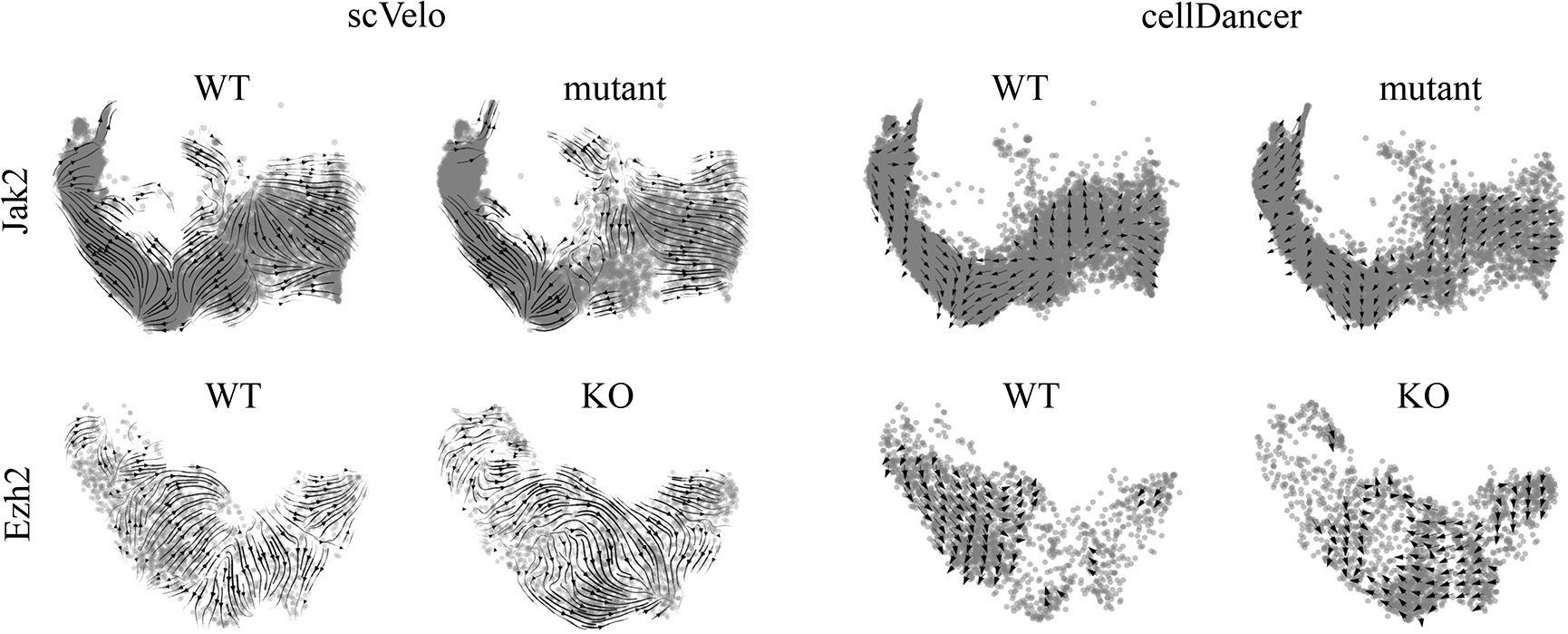
Performance of scVelo and cellDancer. Trajectories recovered by scVelo (left panels) and cellDancer (right panels), for the WT (left subpanels) and KO (right subpanels) conditions in the Jak2 mouse (top row) and the Ezh2 mouse (bottom row).

